# Effect of Foraging and Nest Defense tradeoffs on the Reproductive Success of Wood Storks (*Mycteria americana*)

**DOI:** 10.1101/592840

**Authors:** Alexis Bruant, Simona Picardi, Peter Frederick, Mathieu Basille

**Affiliations:** Department of Wildlife Ecology and Conservation, Fort Lauderdale Research and Education Center, University of Florida, Davie, FL. 33314, USA; Department of Wildlife Ecology and Conservation, P.O. Box 110430, University of Florida, Gainesville, FL. 32611, USA

**Keywords:** foraging trips, intra-specific aggressions, nest attendance, nest takeover, parental care

## Abstract

In many species of birds, parental care is provided by both parents to maximize offspring survival, and there may be important trade-offs between maximizing food gathering and nest protection during the nesting period. The role of parental care in determining reproductive success was investigated in Wood Storks (*Mycteria americana*), and specifically how the trade-off between frequency and duration of foraging trips, and nest protection has contributed to the nesting outcome. Parental behavior of 85 pairs of Wood Storks was monitored throughout the nesting season in two breeding colonies in Palm Beach County, Florida. Wood Storks have gradually increased the frequency, but not the duration, of foraging trips as chicks developed. The ratio of hatchlings to fledglings was positively associated with the frequency of foraging trips during late chick development. Intra-specific aggressions resulting in nest takeovers have affected 32 % of the nests under study. Occurrence of nest takeovers have been higher for later-breeding pairs, and was happened primarily in the first few weeks of incubation, but was not affected by the degree of joint nest attendance of both parents. These results establish a functional link between parental effort and reproductive outcome in Wood Storks, and highlight the importance of frequent foraging trips, but not nest attendance, by parents.

---

Any pre- or post-breeding investment made by a parent that increases offspring survival (parental care; Trivers 1972, Westneat and Sherman 1993, Royle *et al.* 2012), may affect reproductive success (Eggert *et al*. 1998). For many species of birds, including most waterbirds (Del Hoyo *et al*. 1992), biparental care is the rule (> 90%; Silver *et al.* 1985, Cockburn 2006, Harrison *et al.* 2009). The degree of parental care received by offspring can be crucial for their survival and, as a consequence, can affect the reproductive success of the parents (Elowe and Dodge 1989, Dijkstra *et al*. 1990, Boland *et al*. 1997). For instance, quality of parental care (body condition of parents), instead of quality of egg (egg size), has been shown to affect chick survival in Short-tailed Shearwaters (*Puffinus tenuirostris*; Meathrel *et al*. 1993).

Wood Stork’s behavior such as incubation, brooding, number of feedings or nest protection has been studied in several studies (Clark 1980, Bryan and Coulter 1991, Bryan *et al*. 2005), but the effect of Wood Stork’s behavior on reproductive success has not been investigated yet. To our knowledge, no study has quantified the direct impact of food provisioning on survival of chicks in Wood Stork. In particular, altricial and semi-altricial hatchlings cannot feed themselves, yet require a copious and steady flow of nutrients to fuel rapid growth and development (Schwagmeyer and Mock 2008). However, provisioning takes time and energy, and parents should trade off optimal levels of offspring provisioning versus nest defense (Mutzel *et al*. 2013), clutch size (Dijkstra *et al*. 1990), clutch mass (Hébert and Barclay 1988), or parental body condition (Erikstad *et al*. 1997). Moreno *et al*. (1999) showed that increasing the intake of food positively affected reproductive success, but little is known about how the trade-off between food provisioning and nest protection impacts reproductive success.

In Wood Storks (*Mycteria americana*), both parents provide parental care (Kahl 1962, 1971, US Fish and Wildlife Service 2001). Earlier studies suggest that daily care from both parents is strictly necessary to ensure survival of eggs and offspring in this species (Clark 1980, Bryan *et al*. 2005). Survival of nestling Wood Storks is affected by many factors, including predation (Rodgers 1987), human disturbance (Bouton *et al.* 2005), intraspecific aggression that can lead to nest takeover (Bryan and Coulter 1991), contamination with toxic chemicals (Fleming *et al.* 1984, Burger *et al.* 1993), as well as weather conditions, such as storms associated with strong winds (Coulter and Bryan 1995, Bouton *et al*. 2005, Bryan and Robinette 2008). However, the reproductive success of Wood Stork seems to be primarily related to prey availability in the environment (Ogden 1994, Griffin *et al.* 2008), which affects the ability of parents to provide sufficient food to sustain the development of chicks until fledging. Wood Storks feed mainly on fish (Kahl 1962, 1971, Ogden *et al.* 1976), captured in 15–50 cm deep water (Coulter and Bryan 1993) using “tactolocation” (Kahl and Peacock 1963, Kahl 1964, Clark 1979). This technique is extremely sensitive to variations in fish availability. Sufficient rains prior to the breeding season are required to increase wetland water levels and increase prey population growth, followed by decreased water levels during the nesting phase to concentrate prey and ensure efficient foraging when energetic needs are highest (Kushlan *et al*. 1975, Ogden and Nesbitt 1979, Beerens *et al*. 2015). In contrast, heavy rains during the breeding season can result in colony abandonment due to dispersion of prey (Frederick and Collopy 1989, Ramo and Busto 1992).

Biparental contributions to nest protection and food provisioning by Wood Storks have been confirmed in several studies (Kahl 1962, Coulter et al. 1999, Griffin et al. 2008). Wood Stork parents must budget their time efficiently to provide adequate food and at the same time protect their young in the first weeks after hatching, when chicks are unable to defend themselves or to thermoregulate autonomously (Bryan and Coulter 1987). In this first phase, the continuous presence of a parent on the nest is necessary (Clark 1980, Bryan *et al*. 2005). After three weeks, chicks exhibit a behavioral change, becoming aggressive to any approach (con-specifics and other species, Kahl 1971) and are able to thermoregulate (Clark 1980). This increased autonomy allows both parents to forage simultaneously when food requirements of the chicks are at their peak, leaving the nest unattended (Kahl 1962). Nestling Wood Storks remain in nests for 50 to 60 days before fledging (Kahl 1971, Coulter *et al.* 1999) and continue to return to nests to be fed by their parents for another one to three weeks (Kahl 1971, Borkhataria *et al.* 2012).

The trade-off between nest attendance and food provisioning has been noted in many species of birds (Komdeur and Kats 1999, Fontaine and Martin 2006, Tilgar *et al*. 2010). A key assumption in this trade-off is that nest attendance is related to nest success, through several mechanisms, such as direct care for chicks, and defense of nest against predators and conspecifics (Giese 1996, Schmidt and Whelan 2005). To our knowledge, no study has investigated the relationship between parental care (measured as frequency of foraging trips, their duration, and nest attendance) and reproductive success in Wood Storks but has been in other wading birds (Miller and Burger 1978); likewise, the importance of parental attendance during the nestling period has been little studied in this group. Thus, our objective was to determine if parental care (nest attendance and foraging behavior) varies during the nesting season and, if so, if such variation affects reproductive success. We predicted that the number of parental foraging trips would increase with the age of chicks, and the mean duration of foraging trips would decrease.

We also predicted a positive relationship between number of foraging trips and reproductive success. Second, we assessed a possible relationship between the time spent by parents at the nest and the occurrence of takeover events. We predicted that the percentage of time spent at the nest simultaneously by both parents would reduce the likelihood of takeovers.

## METHODS

### Study areas

We collected data on parental activities of 85 pairs of Wood Storks on 61 nests (due to takeovers and abandoned nests) in two colonies in Palm Beach County, Florida. We observed 32 nests on two islands located within the Wakodahatchee Wetlands (26°47’87”N, 80°14’34”W), nesting on pond apple (*Annona glabra*), dahoon holly (*Ilex cassine*), and sabal palm (*Sabal palmetto*) (Bays *et al*. 2000). We also monitored 29 nests in pond apple and sabal palm in the Ballenisles Country Club, situated on a single island within a golf club (26°83’01”N, 80°10’91”W), located 46.6-km north of Wakodahatchee Wetlands (Fig. 1). It is important to note that Palm Beach County and the large adjacent protected marsh lands (including Loxahatchee National Wildlife Refuge) are a hotspot for the year-round distribution of resident wood storks, which may indicate that foraging habitat is generally good in this area (Picardi *et al*. 2020). The 2017 breeding season in Palm Beach County was characterized by lower precipitation (2,68 mm in January and February, 5,26 mm in March through May) than the long-term average (9,33 mm for December through February 1981-2010, 12,76 mm for March-May 1981-2010; data from the National Oceanic and Atmospheric Administration).

**Fig. 1.**
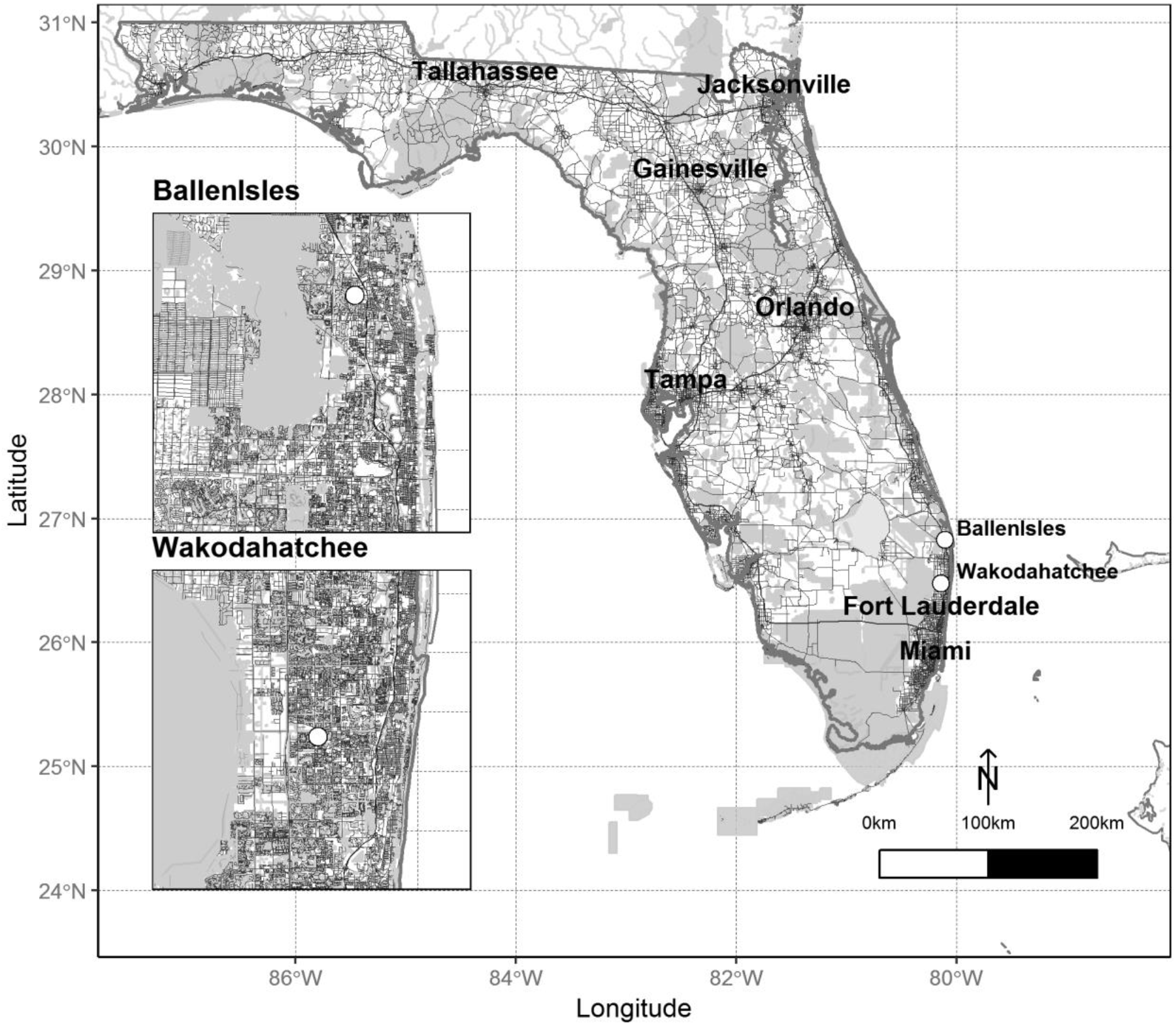
Location of study sites in south Florida, north of Miami and Fort Lauderdale. Protected and natural areas are presented in light gray; roads and interstates as grey and black lines, respectively.

### Data collection

We observed breeding behavior in the two colonies from January 31^st^ to June 2^nd^, 2017. Two observers conducted separate two 5h-long surveys biweekly at each colony, once in the morning (07:30 hr to 12:30 hr) and once in the afternoon (12:30 hr to 17:30 hr), as to homogenize time gaps between successive surveys at a single site (every 3.5 days, Monday morning and Thursday afternoon at BallenIsles, Tuesday morning and Friday afternoon at Wakodahatchee Wetlands). At each site, two groups of nests were observed in weekly alternation by both observers to prevent observer bias. Wood Stork behavior was observed using binoculars (12×50) and recorded using an SLR camera with a 600-mm telephoto lens. We began data collection when most pairs of Wood Storks were either building nests or beginning to incubate eggs. We identified each individual Wood Stork based on unique skin patterns on their head, which are individually unique (Clark 1980, Bryan and Coulter 1991). We built a photographic database for individual identification, consisting of photos of the right and left profile of each stork. Because Wood Storks lack evident sexual dimorphism, sexing partners of a pair was only possible when we witnessed copulation (Clark 1980, Fujioka and Yamagishi 1981, Bryan and Coulter 1991).

At the beginning of each survey, we recorded the status of each nest (construction, incubation or post-hatching, individual parents present), took pictures of the individuals present, and recorded arrivals and departures. Identification was always made later using the photo database. The takeovers were either observed directly or inferred from the data showing the presence of different adults in the nest. A takeover can be carried out by a single individual or by a pair who are seizing an already built nest. This may result in the cessation of egg incubation or the death of the original pairs’ chicks present in the nest at the time of the takeover. Trips were categorized as foraging or gathering nest building material based on whether the parents regurgitated food to nestlings upon return, or brought back twigs and other woody material, respectively. Trips were initially classified as unknown when parents neither fed nor carried nest material when returning. We found that 96% of foraging trips (showing regurgitation) lasted more than 44 min. Using that information, we categorized any unknown trips longer than 44 min as foraging trips. Laying dates for each nest were estimated by back-dating from hatching dates, using an average incubation period of 28 days (Rodgers and Schwikert 1997) and were matched to observations of parental brooding behavior (especially sitting position). We counted hatchlings in each nest where visual counting was feasible. We estimated reproductive success at each nest as the proportion of hatching chicks still alive after 8 weeks, i.e. the estimated time when young birds leave the nest for the first time (Middleton and Prigoda 2001, Bryan *et al*. 2005). We determined early and late pairs according to their nesting date. Most pairs (48 out of 61) initiated nesting within the first week of study, before February 8, 2017, and were considered early nesters. A second wave initiated nesting between February 20, 2017, and April 21, 2017 and were considered late nesters.

### Analytical Methods

#### Changes in frequency and duration of foraging trips

We modeled both frequency and duration of foraging trips as non-linear functions of weeks since hatching using generalized linear mixed models (Mirman 2014). The overall shape of the curve was captured with inclusion of orthogonal polynomials on time up to the fourth order, with individual-within-nest random effects on all terms (for frequency only; for the duration analysis, the use of complete trips with known time of departure and arrival limited the sample size, and the estimation of random effects was not possible). Using a subset of individuals of known sex, sex differences were tested with the inclusion of an additive and multiplicative fixed effect of sex. Similarly, we included an additive and multiplicative fixed effect of the calendar date of nest initiating to test its effect on frequency and duration of foraging trip.

#### Effect of frequency of foraging trips on reproductive success ratio

The effect of frequency of foraging trips on reproductive success was then assessed using a logistic regression on the number of successes (fledglings) over failures (hatchlings that did not survive until fledging) in each nest. Fixed effects of the frequency of foraging trips during the early pre-flight stage (weeks 1–4) and during the late pre-flight stage (weeks 5–8) were included in the regression, after checking for their correlation. We have not reported results for the post-flight stage because the data were not sufficient to constitute a robust data set. In fact, at the end of the young’s development (between 9 and 12 weeks), parents return very little to the nest to feed them. It happened several times that the parents did not return to the nest during the 5.5 hours of observation.

#### Nest attendance and risk of takeover

We fit semi-parametric proportional hazards (SPPH) models to the time spent by parents at the nest prior to takeovers, expressed either as a function of calendar time, or time within the nest cycle. The instantaneous risk of successful takeover of a nest at a given week was modeled as a function of the baseline hazard experienced by all individuals, and the proportion of time with at least one adult or two adults present at the nest during the week. In the semi-parametric approach, weak assumptions about the baseline hazard are made, allowing the estimation of the relative risk of takeover. Covariate effects are then estimated using a partial likelihood that does not require estimating the baseline hazard.

All statistical analyses were performed in the software R 3.3.0 (R Core Team 2017) using notably the packages “lme4” (version 1.1.13; Bates *et al.* 2015), “survival” (version 2.40.1; Therneau and Grambsch 2000), and “cowplot” for graphs (version 0.7.0; Wilke 2016).

## RESULTS

### Wood Stork monitoring

Observations from the two sites were similar. For instance, comparing the two sites we obtained an average foraging rate of 0.243 h^-1^ for Ballenisles Country Club versus 0.239 h^-1^ for Wakodahatchee Wetlands. Moreover, the pairs with the highest chick survival were those with the highest mean frequency of foraging trips per hour in each site. We thus pooled both colonies to increase robustness of our results. Between January 31^st^ and June 2^nd^ 2017, we monitored 61 nests (29 in BallenIsles, and 32 in Wakodahatchee Wetlands), corresponding to 85 nesting attempts from individually identifiable pair of Wood Stork (with sex identified for individuals of 71 pairs). Of these nest attempts, 27 were taken over by another pair, five were abandoned, and 53 either succeeded or were still active at the end of data collection (Fig. 2). Among surviving nests, we were able to track the fate of chicks until fledging for 29 nest attempts (see Table 1). For the other nest attemps, we were not able to track the fate of chicks until fledging due to either takeover, death of chicks, abandoned nest or the end of the observation period. We found an average of 3.00 ± 0.46 SD hatchlings per nest, and an average of 2.59 ± 0.57 SD chicks fledged per nest.

**Fig. 2.**
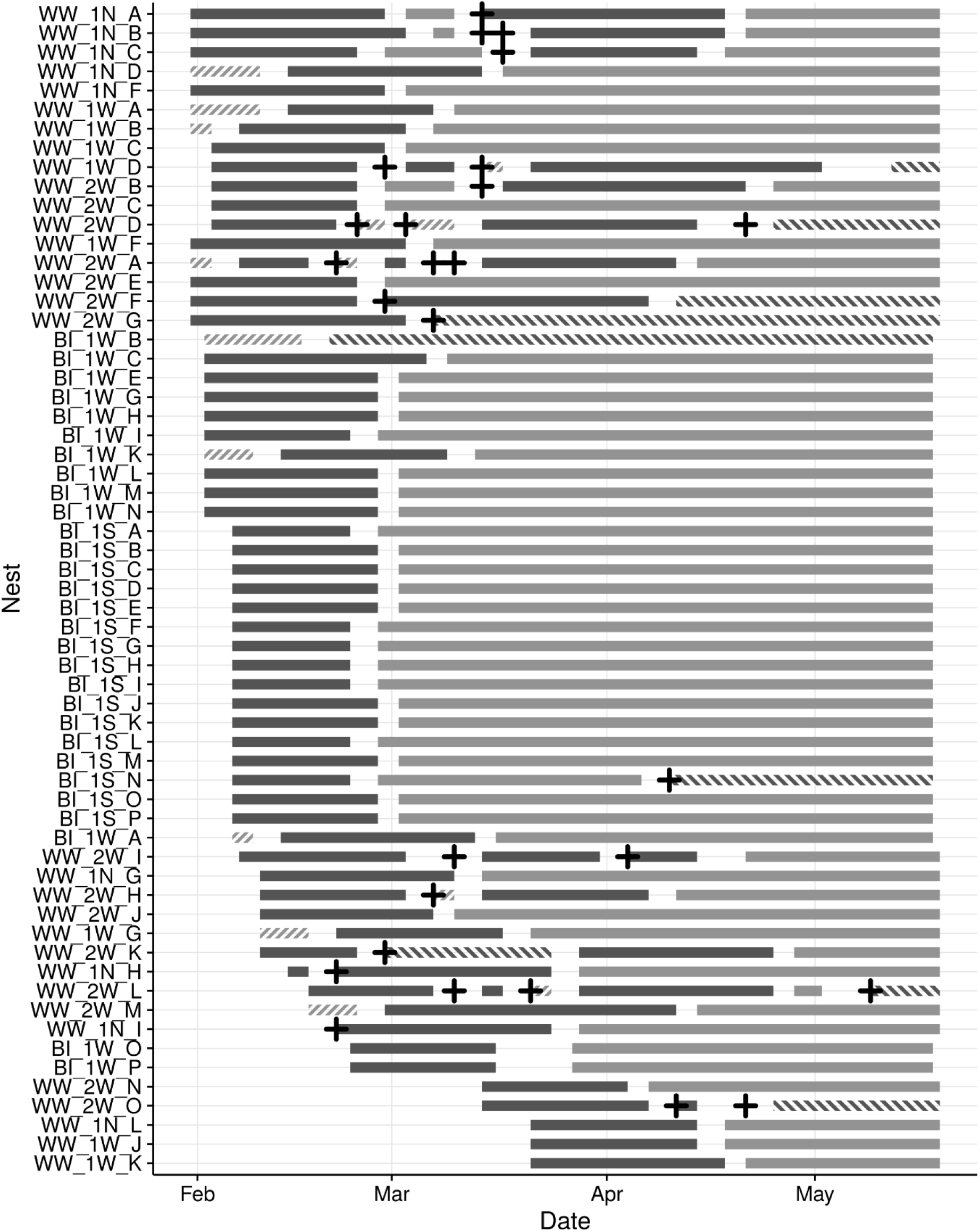
Evolution of nest status during the nesting season in Wood Stork (*Mycteria Americana*) (*n* = 61). The color indicates the phases of each pair in the nest: Building phase (stripped light gray), Incubation phase (dark gray), Post-hatching phase (light gray), Abandon (stripped dark gray). All 27 takeovers are indicated by black crosses.

**Table 1.**
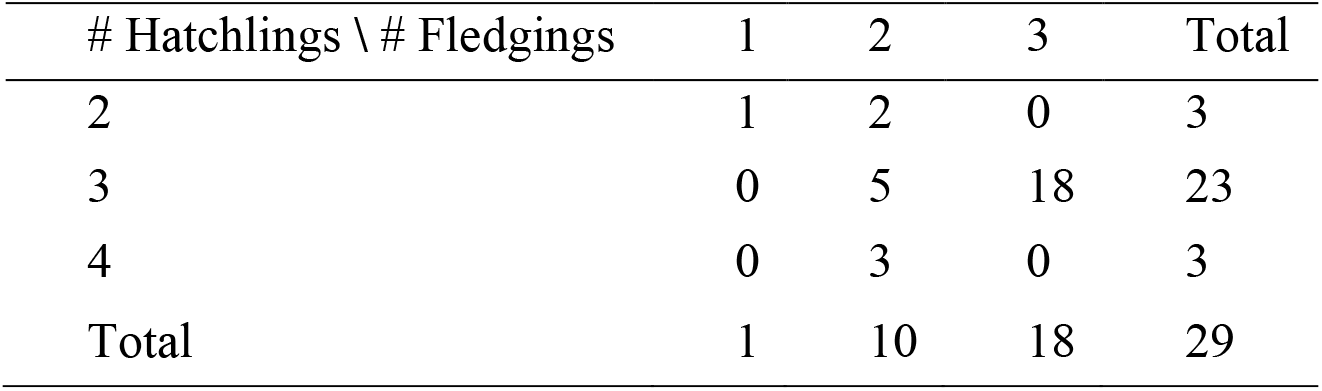
Reproductive success for 29 Wood Stork (*Mycteria Americana*) nests with known outcome in south Florida: Number of nests broken down by the number of chicks per nest at the time of hatching (rows) and at the time of fledging (columns).

### Changes in frequency and duration of foraging trips

Adding orthogonal polynomials successively to the constant model of frequency of foraging trip significantly improved the fit until the quadratic term (χ^2^(7) = 14.792, *P* = 0.039; Table 2A), whereas adding a cubic or quartic term did not further improve the fit further (resp. χ^2^(1) = 1.011, *P* = 0.315; χ^2^(6) = 0.607, *P* = 0.436; Table 2A). Using the quadratic model as a baseline, model selection reaeled that an additive or multiplicative effect either sex (Table 2B) or initiation date (Table 2C) did not significantly improve the fit (Table 2B). The model including the effect of first- and second-order polynomials was kept for the rest of analyses (Table 3). This model showed a significant effect of the first-order orthogonal polynomial term (0.225 ± 0.06, *t*910 = 13.501, *P* < 0.001; Table 3) demonstrating a positive linear relationship between the frequency of foraging trips and the progression of chick development (Fig. 3A).

**Fig. 3.**
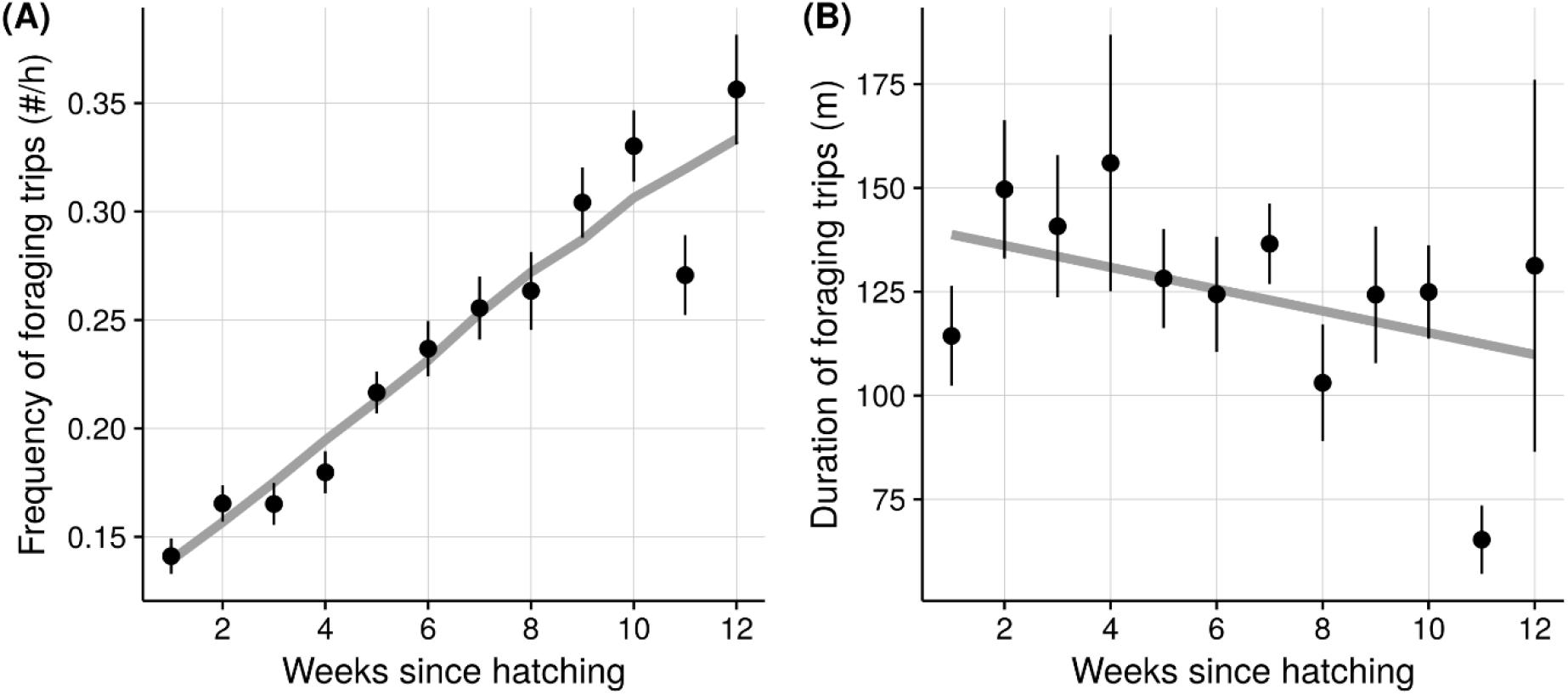
Frequency (A) and duration (B) of foraging trips through time during the posthatching phase in Wood Stork (*Mycteria Americana*) (*n* = 157, *r*^2^ = 0,22 and *r*^2^ = 0,02 respectively). Point represent average values for each week since hatching ± SE, and the light gray line represents the best model fit (see text for details).

**Table 2.** Model selection for frequency of foraging trips by Wood Storks (*Mycteria Americana*). *K* is the number of parameters in the model; AIC the Akaike Information Criterion, LL the log-likelihood, and χ^2^, df and Pr(>χ^2^) indicates the statistic, the associated degrees of freedom and *p*-value for the comparison between each model and the previous one. The selected model is indicated in bold.

| *General shape* |  |  |  |  |  |  |
| --- | --- | --- | --- | --- | --- | --- |
| Model | *K* | AIC | LL | $\chi^2$ | df | $\text{Pr}(>\chi^2)$ |
| Constant | 4 | -1197.7 | 602.83 |  |  |  |
| First-order polynomial | 9 | -1447.3 | 732.67 | 259.679 | 5 | <0.001 |
| **Second-order polynomials** | **16** | **-1448.1** | **740.07** | **14.792** | **7** | **0.039** |
| Third-order polynomials | 17 | -1447.1 | 740.57 | 1.011 | 1 | 0.315 |
| Fourth-order polynomials | 18 | -1445.8 | 740.87 | 0.607 | 1 | 0.436 |
| *Sex effect* |  |  |  |  |  |  |
| Model | *K* | AIC | LL | $\chi^2$ | df | $\text{Pr}(>\chi^2)$ |
| **Baseline** | **16** | **-611.00** | **321.50** |  |  |  |
| Sex (additive) | 17 | -609.86 | 321.93 | 0.858 | 1 | 0.354 |
| Sex (multiplicative) | 19 | -607.32 | 322.66 | 1.463 | 2 | 0.481 |
| *Effect of the start date of nesting* |  |  |  |  |  |  |
| Model | *K* | AIC | LL | $\chi^2$ | df | $\text{Pr}(>\chi^2)$ |
| **Baseline** | **16** | **-1448.1** | **740.07** |  |  |  |
| Start (additive) | 17 | -1446.7 | 740.35 | 0.576 | 1 | 0.448 |
| Start (multiplicative) | 19 | -1448.2 | 743.10 | 5.503 | 2 | 0.064 |

**Table 3.** Coefficients and their significance of the best model for the frequency of foraging trips during the post-hatching phase in Wood Stork (*Mycteria Americana*).

| Variable | Estimate | Std Error | df | *t* | Pr(>\|*t*\|) |
| --- | --- | --- | --- | --- | --- |
| Intercept | 0.225 | 0.006 | 47.1200 | 38.064 | <0.001 |
| First-order polynomial | 1.887 | 0.1403 | 42.630 | 13.501 | <0.001 |
| Second-order polynomial | 0.034 | 0.128 | 67.220 | 0.265 | 0.792 |

Adding orthogonal polynomials to the constant model of duration of foraging trip did not significantly improve the fit (all *P* > 0.05; Table 4A), although the first-order polynomial was close to statistical significance (*F*_1,155_ = 3.612, *P* = 0.059; Table 4A). We thus simplified the baseline foraging duration model as a simple linear model including a main effect of the number of weeks since hatching. Using the simple linear model as a baseline, model selection then showed that the additive or multiplicative effect of sex (Table 4B) and initiation date (Table 4C) did not significantly improve the fit. The simple linear model including no effect of sex or start time of nesting was thus kept for the rest of analyses, and indicated a weak trend of decreasing duration of foraging trips through time since hatching (−2.629 ± 1.388, *t*_27_ = −1.893, *P* = 0.060; Table 5; Fig. 3B).

**Table 4.** Model selection for duration of foraging trips by Wood Storks (*Mycteria Americana*). SSR and SSE are the residual and explained sums of squares, respectively with their associated degrees of freedom, and *F*, and Pr(>*F*) indicate the statistic and *P*-value for the comparison between each model and the previous one. The selected model is indicated in bold.

| *General shape* |  |  |  |  |  |  |
| --- | --- | --- | --- | --- | --- | --- |
| Model | SS<sub>R</sub> | df | SS<sub>E</sub> | df | $F$ | $\text{Pr}(>F)$ |
| **Constant** | **502825** | **156** |  |  |  |  |
| First-order polynomial | 491459 | 155 | 11366.4 | 1 | 3.612 | 0.059 |
| Second-order polynomials | 487172 | 154 | 4286.4 | 1 | 1.362 | 0.245 |
| Third-order polynomials | 481410 | 153 | 5762.2 | 1 | 1.831 | 0.178 |
| Fourth-order polynomials | 478286 | 152 | 3124.5 | 1 | 0.993 | 0.320 |

| *Sex effect* |  |  |  |  |  |  |
| --- | --- | --- | --- | --- | --- | --- |
| Model | SS<sub>R</sub> | df | SS<sub>E</sub> | df | $F$ | $\text{Pr}(>F)$ |
| **Baseline** | **236086** | **69** |  |  |  |  |
| Sex (additive) | 234611 | 68 | 1475.0 | 1 | 0.427 | 0.516 |
| Sex (multiplicative) | 231545 | 67 | 3066.5 | 1 | 0.887 | 0.350 |

| *Effect of the start date of nesting* |  |  |  |  |  |  |
| --- | --- | --- | --- | --- | --- | --- |
| Model | SS<sub>R</sub> | df | SS<sub>E</sub> | df | $F$ | $\text{Pr}(>F)$ |
| **Baseline** | **491459** | **155** |  |  |  |  |
| Start (additive) | 488720 | 154 | 2738.7 | 1 | 0.872 | 0.352 |
| Start (multiplicative) | 480680 | 153 | 8040.3 | 1 | 2.559 | 0.112 |

**Table 5.** Coefficients and their significance of the best model for the duration of foraging trips during the post-hatching phase in Wood Stork (*Mycteria Americana*).

| Variable | Estimate | Std Error | *t* | Pr(>\|*t*\|) |
| --- | --- | --- | --- | --- |
| Intercept | 141.384 | 9.442 | 14.974 | <0.001 |
| Weeks | -2.629 | 1.388 | -1.893 | 0.060 |

### Effect of frequency of foraging trips on reproductive success

We divided the post-hatching phase into three stages: early pre-flight (weeks 1–4 posthatching), late pre-flight (weeks 5–8) and post-flight (weeks 9–12; Clark 1980). The frequency of foraging trips increased during each stage of chick development. However, the effect of the frequency of foraging trips on reproductive success was not significant during the early preflight stage (weeks 1–4; *Z* = 1.025; *P* = 0.306; Table 6). Frequency of foraging trips during the late pre-flight stage had a significant positive effect on reproductive success (*Z* = 2.929; *P* = 0.003; Table 6; Fig. 4). The frequency of foraging trips had no effect on the absolute number of fledglings (*Z* = 1.035; *P* = 0. 438). Pairs with the highest chick survival (*P* = 1) were those with the highest mean frequency of foraging trips per hour (0.25 to 0.35 h^-1^), whereas pairs with the lowest chick survival (*P* = 0.5) had the lowest mean frequency of foraging trips per hour (0.12 to 0.16 h^-1^). The mean frequency of foraging trips during the early and late pre-flight stages were not correlated (*r* = 0.216, t_27_ = 1.148, *P* = 0.261).

**Fig. 4.**
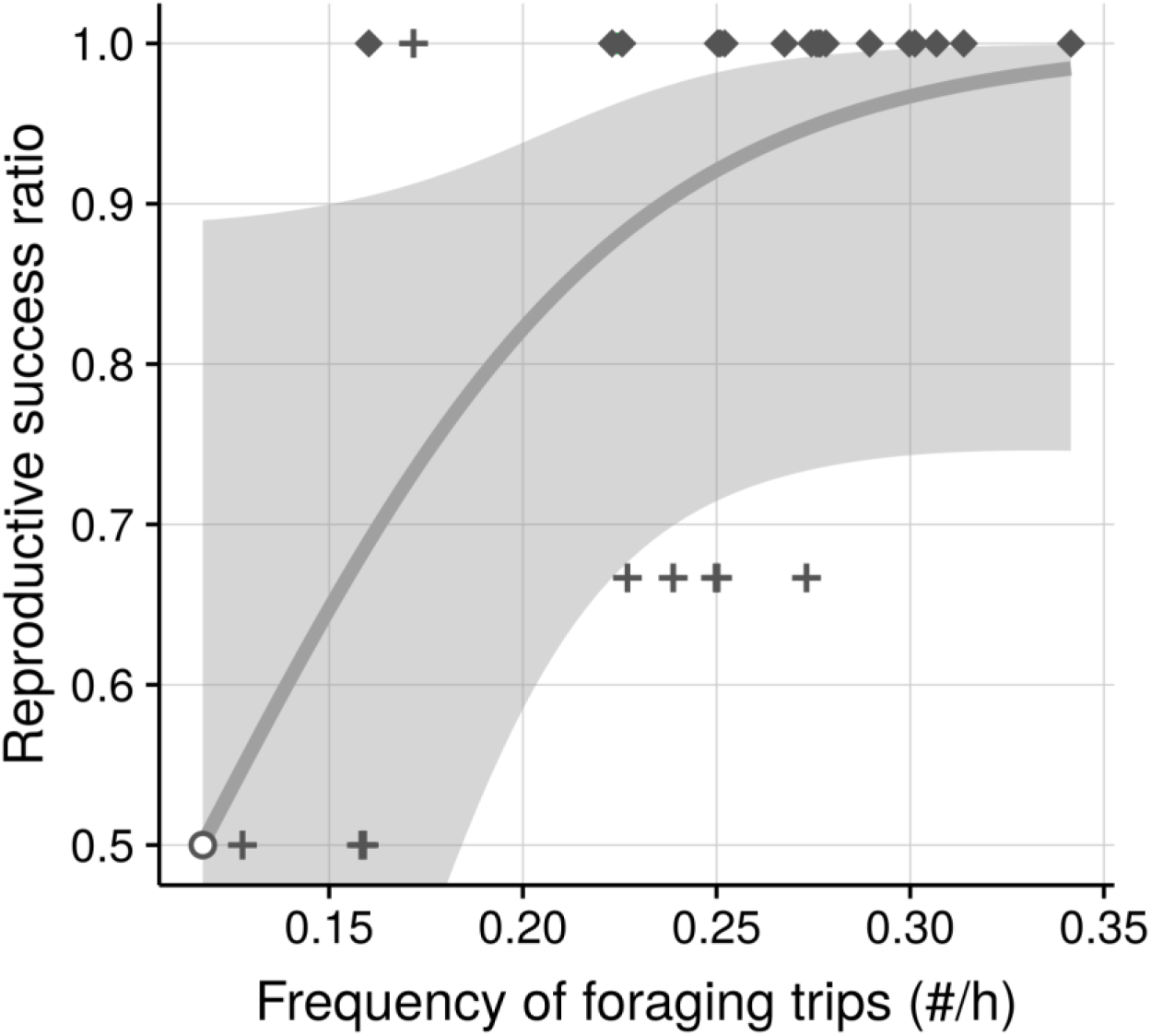
Reproductive success ratio as a function of frequency of foraging trips in late preflight stage in Wood Stork (*Mycteria Americana*) (*n* = 29, *r*^2^ = 0,43). The color of dots indicates the number of fledglings in each nest (diamonds = 3; crosses = 2; circle = 1) and the blue line indicates the logistic fit (with 95 % confidence interval).

**Table 6.**
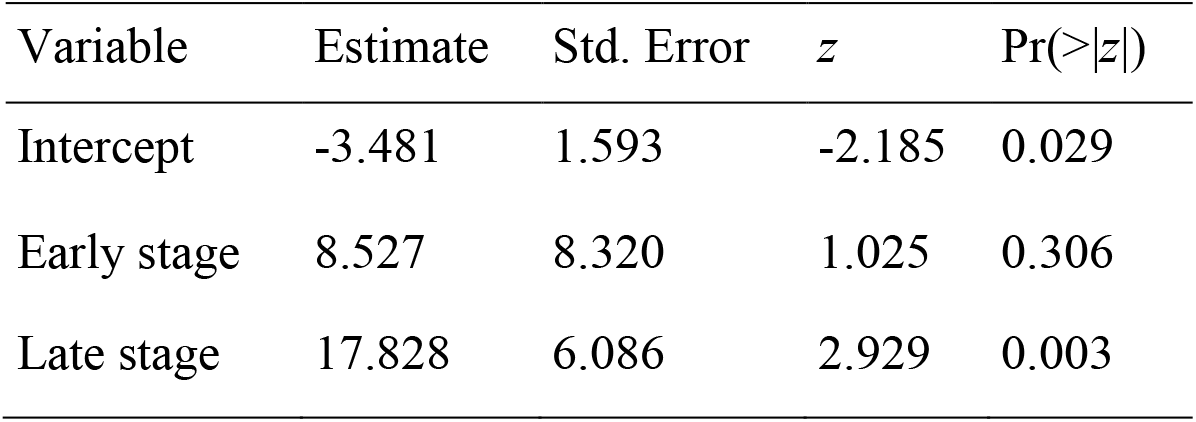
Coefficients and their significance of the logistic model for reproductive success ratio in Wood Stork (*Mycteria Americana*).

| Variable | Estimate | Std. Error | $z$ | $\text{Pr}( > \vert z\vert )$ |
| --- | --- | --- | --- | --- |
| Intercept | -3.481 | 1.593 | -2.185 | 0.029 |
| Early stage | 8.527 | 8.320 | 1.025 | 0.306 |
| Late stage | 17.828 | 6.086 | 2.929 | 0.003 |

**Table 7.** Coefficients and their significance of the semi-parametric proportional hazards (SPPH) model applied to the risk of takeovers of Wood Stork (*Mycteria Americana*) nests. Exponentiated coefficients can be interpreted as multiplicative effects on the hazard, i.e. the instantaneous risk of takeover, holding other covariates constant. For instance, late starters have a risk more than twice as high as early starters (*e^β^* = 2.135).

| Variable | $\beta$ | $e^\beta$ | Std. Error | $z$ | $\Pr(>\vert z\vert )$ |
| --- | --- | --- | --- | --- | --- |
| Presence $\geq 1$ ad. | 0.029 | 1.030 | 0.023 | 1.289 | 0.197 |
| Presence 2 ad. | 0.017 | 1.018 | 0.007 | 2.506 | 0.012 |
| Late-starters | 0.759 | 2.135 | 0.385 | 1.971 | 0.049 |

### Nest attendance and risk of takeover

Both adults of each pair were simultaneously present at nests more than half of the time during nest building and the first week of incubation, but, this proportion declined rapidly during incubation (Fig. 5A). Until the beginning of the early pre-flight stage, at least one adult was always constantly at the nest. Then, the presence of even one adult gradually decreased, reaching a minimum of 10% of the time at the end of the nesting season (Fig. 5A). In our study, 32% of nests experienced a takeover. Of the 27 takeovers, the vast majority took place in Wakodahatchee Wetlands (26) vs. only 1 in Ballen Isles. Most takeovers occurred between February 13 to March 19 2017 (Fig. 5B). However, we found that the amount of time with presence of at least one parent at the nest did not significantly affect risk of successful takeovers (*e^β^* = 1.030, Z = 1.289; *P* = 0.197; Table 7), whereas a higher presence of both parents at the nest had a significant positive effect on the risk of takeover (*e^β^* = 1.017; Z = 2.506; *P* = 0.012; Table 7). Similarly, the date of initiation of incubation had a positive effect on the risk of takeovers, with later initiation dates associated with higher risk (*e^β^* = 2.135; Z = 1.971; *P* = 0.048; Table 7; Fig. 5C).

**Fig. 5.**
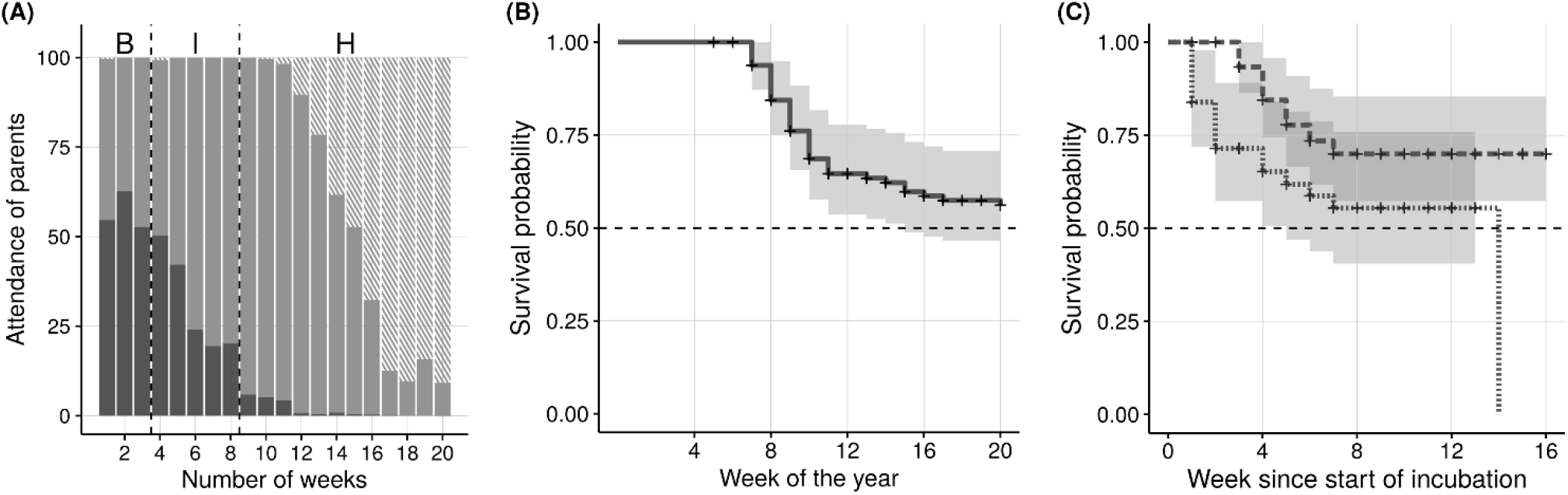
(A) Nest attendance by adult Wood Storks (*Mycteria Americana*) through time (*n* = 79 pairs). The color indicates nest attendance of two adults (dark gray), one adult (light gray) or no adults (stripped light gray). B: building phase; I: incubation phase; H: posthatching phase. The average fledge age is between 50 to 60 days (Kahl 1971, Coulter *et al.* 1999, Bryan *et al*. 2005). (B) Kaplan-Meier survival curve (with confidence interval) for the risk of nest takeovers in 79 Wood Stork nests as a function of the week of the year (*n* = 79 pairs), or (C) week since start of incubation (*n* = 79 pairs). Colors further distinguish between early pairs (dashed lined, top), and late pairs (dotted line, bottom).

## DISCUSSION

We found that the mean frequency of foraging trips per hour showed a gradual increase in time with the developmental stage of chicks. However, our sample of chicks in the postflight stage was small and highly variable among different pairs, thus limiting our inference during the final stage. The duration of foraging trips was not influenced by the developmental stage of chicks. These results are in general agreement with those of previous studies of Wood Storks (Clark 1980, Bryan *et al.* 1995, 2005). Parents meet the increased energetic needs of chicks (Kahl 1962) by increasing the number rather than the duration of foraging trips (Bryan *et al.* 1995). During the post-hatching phase, the mean frequency of foraging trips per hour and their duration were not different between the sexes, and were not related to the onset date of incubation. Whether or not the duration of foraging trips is always stable as chicks grow is unclear, but may depend on the mosaic of wetland conditions in the area surrounding at colony (Coulter and Bryan 1993, Bryan *et al.* 1995). The dry season was not interrupted by major rainfall events that could reverse gradual water drydown, and this resulted in good conditions for wood stork foraging in our study area (Kushlan 1986). However, precipitation in summer 2016 were lower (7,7 mm) than the long-term average (22,01 mm) in Palm Beach County. High water levels in the prior non-breeding season promote growth of fish populations (DeAngelis et al. 2010, Botson et al. 2016), so it is possible that the low levels observed in 2016 prevented foraging conditions from reaching optimality in 2017, despite the favorable trends of water recession during the breeding season.

We found that the mean rate of foraging trips in the late pre-flight stage, but not in the early pre-flight stage, affected the proportion of chicks that fledged. Although our results showed that pairs with the highest chick survival were those with the highest mean frequency of foraging trips per hour and that pairs with the lowest chick survival had the lowest mean frequency of foraging trips per hour, it is important to note the possible circularity of this relationship: in fact, one could argue that pairs that reached the late pre-flight stage with an already reduced brood size consequently decreased the frequency of trips due to reduced demands from the offspring, rather than the other way around. However, our results show that the relationship between reproductive success and frequency of foraging trips is independent of the absolute number of fledglings. For example, pairs with two successful chicks but low fledging success had either higher (0.23–0.28 h^-1^ for success ratio of 0.67, see Fig. 2) or lower (0.13–0.16 h^-1^ for success ratio of 0.5, see Fig. 4) foraging trip frequency than those with 100 % success ratio (0.17–0.23 h^-1^, see Fig. 4). If the absolute number of chicks was driving the frequency of foraging trips, we would expect the same frequency for an equal number of chicks independently from the initial brood size. This result supports an effect of foraging trip frequency in determining chick survival ratio, and not vice-versa. Because we found no effect of the frequency of foraging trips on the absolute number of fledglings, fledging success is apparently determined by other factors, such as mortality related to predation, sibling competition, or disturbance, as well as factors affecting clutch size (Burger 1982, Rodgers 1987, Bouton *et al.* 2005).

Takeover behavior appears to be widespread in Wood Storks, where it may affect more than a third of pairs in a colony (Bryan and Coulter 1991). In our study, 32% of nests experienced a takeover. The risk of a takeover was more than double for late pairs (nest initiation after February 13^th^). The lower occurrence of takeovers for early pairs could be explained by better intrinsic characteristics of individuals (Johnson and Kermott 1990) such as larger size, higher energy reserves, greater aggressiveness, or higher social status (i.e. dominance hierarchy rank). We also found that the time of simultaneous nest attendance by both parents did not reduce the occurrence of takeovers. On the contrary, and surprisingly, pairs with greater nest attendance times by both individuals had a greater chance of undergoing a takeover. Because Bryan and Coulter (1991) found that all takeovers occurred when a single individual was present at a nest, we expected that pairs that spent more time together at the nest would have a lower chance of getting their nest taken over, but, instead, we found that the presence of both individuals was associated with a greater risk of undergoing a takeover. Pairs may increase joint attendance when their perceived risk of attack is greater, but we have no data to test this possible explanation. In any case, increased parental attendance did not result in reduced risk of nest failure, suggesting that there may not be a clear trade-off between attendance and time spent foraging. Perhaps attendance of both parents at the nest does not provide appropriate protection against takeovers, and body condition of parents may be a determining factor as shown in the study by Meathrel *et al*. (1993) in Short-tailed Shearwaters.

Wood Storks have long breeding cycles (~110 days), and are often limited by food availability at the end of the dry season (Kushlan *et al.* 1975). Our results suggest that the frequency of foraging trips during the late pre-flight stage is positively associated with the reproductive success of Wood Storks. To our knowledge, this is the first time a link is established between parental effort and reproductive success for this species. Moreover, our findings highlighted an important mechanism of nest failure, nest takeovers, and that the risk associated with it varies throughout the season. Thus, in addition to constraints related to seasonal water level fluctuations, a late nesting initiation can be associated to low reproductive success due to a higher probability of takeover events. It is important to point out that we have one season of data and that the reproductive success of Wood Stork is primarily related to prey availability in the environment (Ogden 1994, Griffin *et al.* 2008). This food availability is important for the development of chicks until fledging. Thus, a study carried out over several breeding seasons in the same location could provide more accurate breeding estimates as foraging conditions are a dynamic system component.

## ACKNOWLEDGMENTS

We are grateful to Gary Myers and the management office of BallenIsles Country Club and to Robert Nelton and the Palm Beach County Water Utilities Department for granting us access to their premises. Gary Ritchison provided valuable feedback on an earlier version of this manuscript. This work was supported by the USDA National Institute of Food and Agriculture (Hatch project 1009101).

## Supplemental Material

**Plate 1.**
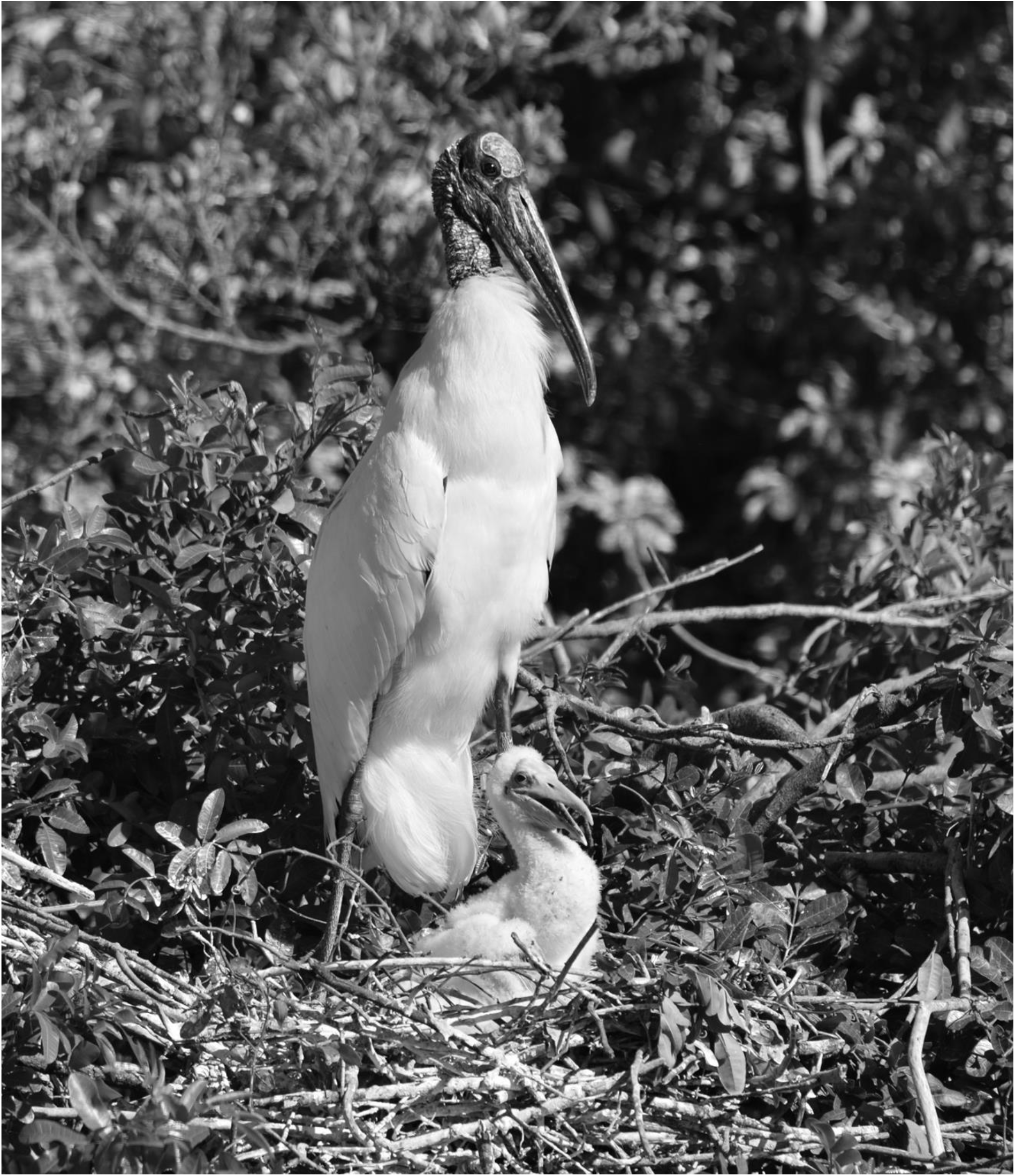
An adult Wood Stork (*Mycteria americana*) attending its nest, with a three-weeks old nestling. Study of nest attendance over time showed it had not effect on the risk of conspecific taking over the nest, and ultimately on nest success.

